# The highly heterozygous European amphioxus (*Branchiostoma lanceolatum*) at the edge of panmixia

**DOI:** 10.64898/2026.01.13.699254

**Authors:** Marina Brasó-Vives, Diego A. Hartasánchez, Angélica Pulido, Stéphanie Bertrand, Hector Escriva, Marc Robinson-Rechavi

## Abstract

Amphioxus (Cephalochordata) are small marine chordates that have broad ecological ranges, yet as adults form local settlements and exhibit limited mobility. Genomic surveys of two amphioxus species have suggested that they rank among the most genetically diverse metazoans. Here, we present the first accurate assessment of genomic diversity in the European amphioxus (*Branchiostoma lanceolatum*) and investigate the processes underlying this diversity. We leverage whole-genome sequencing data from multiple individuals sampled at two geographically distant Atlantic and Mediterranean locations. Consistent with previous estimates in other amphioxus species, we measure exceptionally high genomic diversity, with an average heterozygosity of 2.73% in *B. lanceolatum*. Despite the large geographic separation between sampling sites, population differentiation is minimal, indicating extensive gene flow among distant adult settlements. Phylogenetic analyses combined with population genetic simulations confirm that this elevated genomic diversity is primarily driven by a large effective population size. Although adult amphioxus have limited mobility, our results indicate that long-distance larval dispersal mediated by ocean currents is sufficient to generate a near-panmictic population structure across their broad ecological range.

**Author Summary:** Amphioxus are small marine animals that are close relatives of vertebrates. Although they appear simple and are very similar among species, previous studies have suggested that their genome may be more diverse than expected. In this study, we accurately measured genetic diversity across the entire genome of European amphioxus collected from the Atlantic Ocean and the Mediterranean Sea. We found that this species exhibits a genetic diversity among the highest reported in animals.

Despite being collected from far-apart locations, individuals are genetically very similar. This suggests that amphioxus populations are strongly interconnected. While adults move very little, their larvae drift with ocean currents, enabling them to travel long distances before settling. This widespread mixing helps to maintain a single, large, interconnected population.

Our results demonstrate that even species that appear simple and sedentary can exhibit extraordinary genetic diversity when they have large populations and effective long-distance dispersal mechanisms. Understanding how such diversity is generated and maintained is essential for comprehending evolution and marine biodiversity.

## Introduction

*Genomic diversity* refers to the extent to which the genomes of individuals within a species differ from one another. The greater the species’ genomic diversity, the greater its capability to adapt to environmental changes and, thus, to survive. Species with very low genomic diversity are at a higher risk of extinction [1]. For this reason, the study of genomic diversity and the forces behind it have been of great interest to those studying molecular evolution, ecology, and population genetics for decades [2–9]. While the theoretical determinants of genomic diversity are largely understood, identifying the causes of genomic diversity in specific cases remains challenging [2,3,5,6]. Improving our understanding of the factors that determine the genomic diversity of specific species could help the development of targeted conservation efforts for endangered species [1,10,11]. In this study, we examine the extreme case of amphioxus genomic diversity, which is recognized as one of the highest among metazoans [12–14].

Cephalochordates, also known as lancelets or amphioxus, are a small group of chordates that constitute the sister-group of the Olfactores, which includes vertebrates and tunicates. They are small marine filter-feeders with only short-distance mobility as adults, typically producing local settlements [15,16]. They reproduce through external fertilization and have larval stages that last for long periods of time (weeks to months) and disperse through marine currents [15–19]. Here we study the European amphioxus (*Branchiostoma lanceolatum*), which has a wide ecological distribution, spanning at least all the coasts of the Mediterranean Sea and the northeastern Atlantic Ocean from North Africa to the Baltic Sea [15,16,20].

Estimates of genomic diversity depend heavily on the type of data used and the methodology employed. In 2008, the first genome reference of an amphioxus species, *Branchiostoma floridae,* revealed one of the highest estimates of genomic diversity ever measured for a metazoan species: 4.0% of heterozygosity [12]. Similarly high figures were later estimated for another amphioxus species, *Branchiostoma belcheri*: 5.37% estimated in 2014 [13] and 3.04% estimated in 2020 [14]. The first two estimates were performed on single individuals, with heterozygosity estimates extracted from differences between reads coming from different chromosome pairs [12] or diploid genome assembly sequences [13]. The latter estimate, on *B. belcheri*, used whole-genome sequencing data on a sample of individuals collected from a single sampling site and mapped to a scaffold-level genome assembly [14]. The difference between the two *B. belcheri* estimates is illustrative of how these estimates are dependent on method and sample size. Overall, despite the differences due to methodology, these three independently measured estimates converge on finding that amphioxus species are among the most genomically diverse among metazoan species. They are comparable only to a very few species of molluscs, arthropods, nematodes, and tunicates [5,7,8,21].

While natural selection may be responsible for genomic diversity in a specific region of the genome (e.g., a gene), diversity on a genome-wide scale is mainly governed by forces other than selection [22]. Theoretically, in a population with random mating (i.e., panmictic) and long-term stable population size, heterozygosity should approach the *population-scaled mutation rate* (θ) which is directly proportional to the *effective population size* (N_e_) and the *mutation rate* (μ; θ = 4N_e_μ) [2,23]. In practice, reproductive strategies far from random mating, fluctuations in population size, population substructure, and introgression events can greatly influence genomic diversity [6].

Here, we use European amphioxus medium-coverage whole-genome sequencing data from multiple individuals collected in two distant natural populations (Atlantic and Mediterranean). We map this data to a high-quality assembly, which robustly distinguishes alleles from paralogs [24]. We rigorously measure genomic diversity in all of the individuals, find very little population substructure despite the distant sampling sites, and analyze further possible causes of high genomic diversity. We reach the conclusion that, despite their local settlements and low mobility during late stages, the European amphioxus is at the edge of panmixia and has a very large population size encompassing all of its ecological range.

## Results

We performed whole-genome sequencing of 36 European amphioxus (*Branchiostoma lanceolatum*) adult individuals collected from the wild. Our sampling includes 9 females and 10 males collected in Roscoff (France), on the European Atlantic coast, and 9 females and 8 males collected in Banyuls-sur-mer (France), on the European Mediterranean coast (Figure 1A). We used BWA MEM2 [25] to map the sequencing reads to the *B. lanceolatum* assembly BraLan3 [24], with a resulting mean coverage of 13.26X (Supplementary table 1). In order to accurately call single nucleotide polymorphisms (SNPs) and insertions or deletions (INDELs), we followed the best practices of GATK for non-model organisms [26], including two rounds of base recalibration and a conservative hard filtering (see Methods; Supplementary figure 1; Supplementary table 2). To assess the robustness of our methods we tested an alternative genome mapper, Minimap2 [27], and an alternative variant caller, FreeBayes [28] on chromosomes 16, 17, 18 and 19.

**Figure 1.**
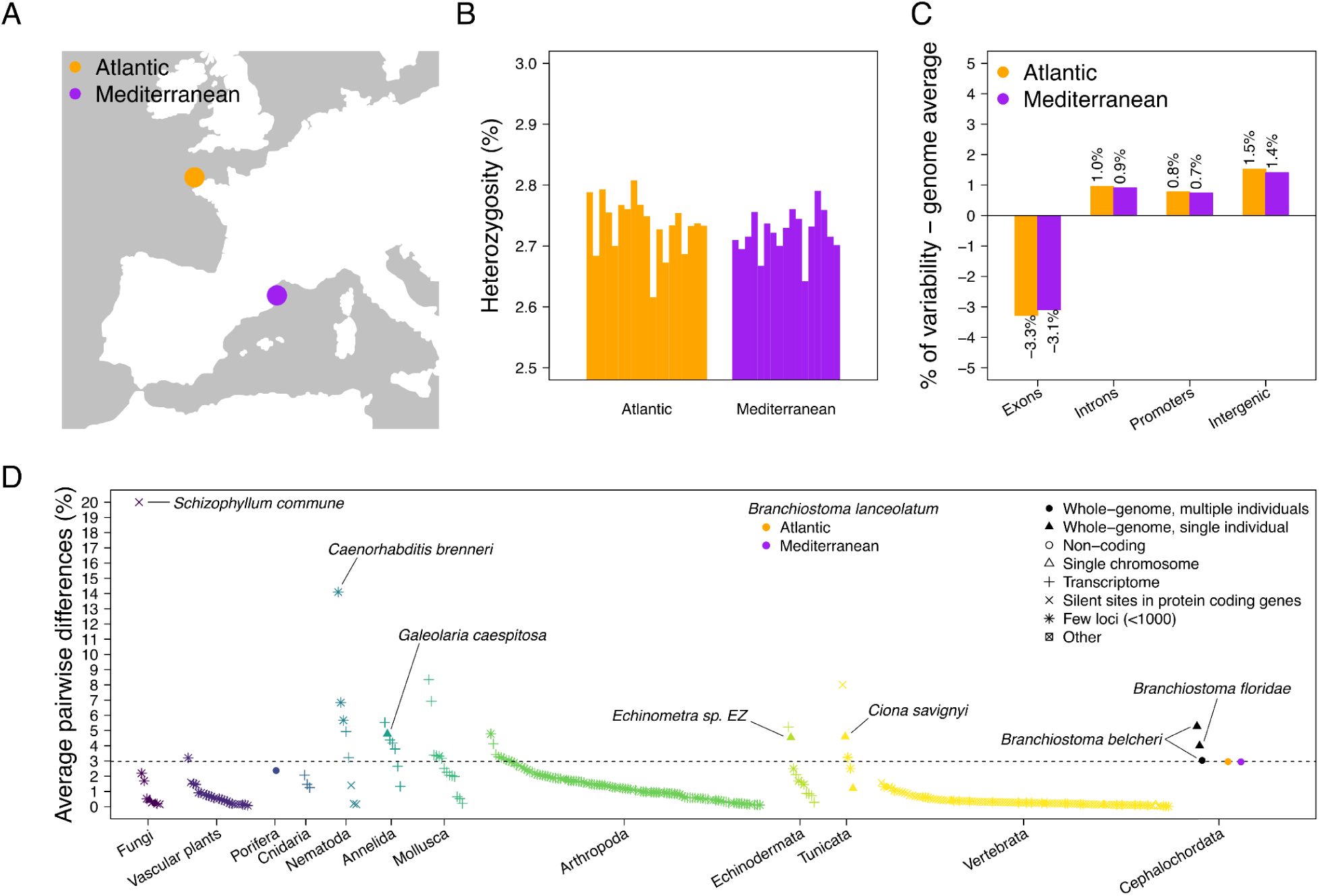
*B. lanceolatum* sampling and genomic diversity. A) Sampling sites for 2 populations of *B. lanceolatum* in Roscoff (Atlantic) and Banyuls-sur-mer (Mediterranean). Image from freesvg.org with Creative Commons 0 license. B) Genomic diversity per individual in both populations, measured in percentage of heterozygous sites. C) Deviation from genome-wide average percentage of variant sites in exons, introns, promoters and intergenic regions for each population. All cases are significantly different from genome-wide average (Fisher’s exact test p-values < 2.2 · 10^-16^ in all cases). D) Across species average pairwise differences (π) comparison. The dashed line corresponds to the average *B. lanceolatum* estimate from this study, while population π estimates are depicted in circles in orange (Atlantic) and purple (Mediterranean) on the right of the figure. A non-exhaustive list of other species estimates are classified by major taxonomic groups (color) and methodology (point shape). Species names of the most relevant cases are specified (either the higher estimates or the top ones performed with whole-genome data). Estimates were extracted from three compilations and several independent studies (Supplementary table 4) [5,8,12–14,30–44].

In addition to the already restrictive variant-calling methods, we only considered positions for which we were able to very reliably call variants with our data in the *B. lanceolatum* genome, hereafter referred to as *callable genome* (see Methods; Supplementary figure 2; Supplementary table 3). To obtain it, we used three filtering criteria (see Methods; Supplementary table 2). First, we limited our analysis to genomic regions with unambiguous genomic mappability, excluding repetitive and duplicated regions, using the software SNPable (see Methods). Second, we only considered positions with a depth of coverage of at least 5X in each one of the samples. Finally, we performed a pruning of isolated variants. As a result, our callable genome was restricted to 40.34% of the *B. lanceolatum* genome (185,241,374 base pairs out of 459,158,900 base pairs of BraLan3), composed of unique mappable regions for which we could call both homozygous and heterozygous variants in all of our samples with high accuracy (see Methods; Supplementary table 2). Callable regions are present along all BraLan3 chromosomes, except for candidate telomeric and centromeric regions (Supplementary figure 2A, 2B, 2C). They are enriched in exons and promoters, and depleted in introns and intergenic regions (Supplementary figure 2D; Supplementary table 3). All subsequent analyses were performed on this callable genome.

### Genomic diversity

We confidently called 42,618,268 variants; 77.75% of SNPs (33,137,498) and 22.25% of INDELs (9,480,770). This means that, despite our conservative variant calling, a striking 23.01% of the *B. lanceolatum* callable genome is variant among our samples. We estimated a mean heterozygosity of 2.74% among the Atlantic population (with a standard deviation of 4.68·10^-4^) and of 2.72% among the Mediterranean population (with a standard deviation of 3.61·10^-4^; Figure 1B; Supplementary table 1). A t-test comparing the mean heterozygosity of the two populations revealed no significant differences, meaning that the combined heterozygosity is a valid estimate. Mean heterozygosity among all samples is 2.73%. Overall per-site nucleotide diversity (π) [29] is 2.971% (2.956% for the Atlantic population and 2.942% for the Mediterranean population). Given our stringent variant filtering criteria, these are lower-bound estimates of the true diversity. Heterozygosity estimates obtained with alternative genome mapping and variant calling methods show slight differences yet overall robustness (Supplementary figure 3).

*B. lanceolatum*’s exons contain fewer variant sites than intergenic parts of the genome (with a Fisher’s exact test p-value < 2.2 · 10^-16^ and an odds ratio of 0.71; Figure 1C). This is expected given that exons are typically under purifying selection. Introns and promoters also showed significantly fewer variants than intergenic regions with less pronounced odds ratios (Fisher’s exact test p-values < 2.2 · 10^-16^ in all cases and odds ratios of 0.97 and 0.95, respectively). These results show that purifying selection in coding regions is a major force determining variant distribution in the *B. lanceolatum* genome.

With 23.01% of sites showing variants, there is a considerable probability that a new mutation will occur in an already segregating site. Accordingly, we observe a non-negligible proportion of non-biallelic variants: 11.99% of SNPs and 58.03% of INDELs (Supplementary figure 4B) have more than two segregating alleles. This observation implies a clear departure from an infinite site model and has major implications for the application of classical population genomics models and statistics to *B. lanceolatum* (see Discussion).

Our estimates confirm that *B. lanceolatum* has one of the highest levels of genomic diversity among multicellular eukaryotes. Figure 1D illustrates our estimates alongside a non-exhaustive list of diversity estimates from various eukaryotic taxonomic groups (Supplementary table 4). Estimates of genomic diversity depend heavily on the data and methodology used. Estimates based on a few loci or only coding regions (e.g., the transcriptome or silent sites in protein-coding genes) provide poor approximations of the genomic diversity across the entire genome. Even estimates performed with whole-genome data depend on the sample size and quality of the genome reference. An estimate based on a single individual’s whole genome is less accurate than one based on multiple individuals’ whole genomes (Figure 1D) [45]. The diversity of the methods in Figure 1D illustrates how poorly comparable they are (see Discussion). Estimates of genomic diversity in amphioxus (Cephalochordata), including our own, are among the highest when using whole-genome data. Our estimate is unique in that it was performed using whole-genome data from multiple individuals collected from two distant sampling sites and mapped to a high-quality, chromosome-level genome reference. This makes it the most reliable estimate to date. Compared to the other amphioxus species estimates [12–14], our results are among the lowest and are similar to those measured by Bi et al. in 2020 [14]. This is expected, given our conservative approach, which leaves room for a higher actual value.

### Population differentiation analysis

Our sampling included two distant locations separated by the natural barrier of the Strait of Gibraltar (Figure 1A). In order to study the *B. lanceolatum* population substructure between the two sampling locations, we measured population differentiation using the *fixation index* (F_ST_) between the two populations [46]. F_ST_ measures the proportion of genetic diversity between individuals from different populations that can be explained by their population of origin. F_ST_ between Atlantic and Mediterranean *B. lanceolatum* population is equal to 0.00429. This value is very small, an indication of almost no population differentiation and high gene flow between populations. It is comparable, for example, to the low population differentiation values of neighboring European human populations [47].

Principal Component Analysis (PCA; Figure 2A) revealed a first principal component (PC1) representing 25.06% of the variance that cannot be explained neither by population nor sex, while it is PC2, explaining only 2.95% of the variance, that represents population differentiation. These results are in line with what we observed with F_ST_, with population differentiation being a small part of the total diversity present in the genome of *B. lanceolatum*. PC3 (2.22%) and PC4 (2.20%) show no sex nor population differentiation (Supplementary figure 5). We obtained similar results performing PCA considering only SNPs, only biallelic variants, discarding singletons and doubletons, or only exonic variants (Supplementary figure 5). PCA using variants stratified by major allele frequency showed that the percentage of the variance explained by PC1 decreases with major allele frequency (Supplementary figure 5). This means that medium-frequency variants, not low-frequency variants, are responsible for the variance captured by PC1. That is, rare variants are more population specific, while variants with higher frequencies tend to be shared among the two populations, also a sign of very low population differentiation and presence of gene flow. PCA performed with alternative genome mapping and variant calling methods show a similar PC1 in all cases (22 to 25% of explained variance and no split of populations or sexes), while PC2 fails to differentiate the two populations in some cases. Overall, PCA results are robust across methods (Supplementary figure 3).

**Figure 2.**
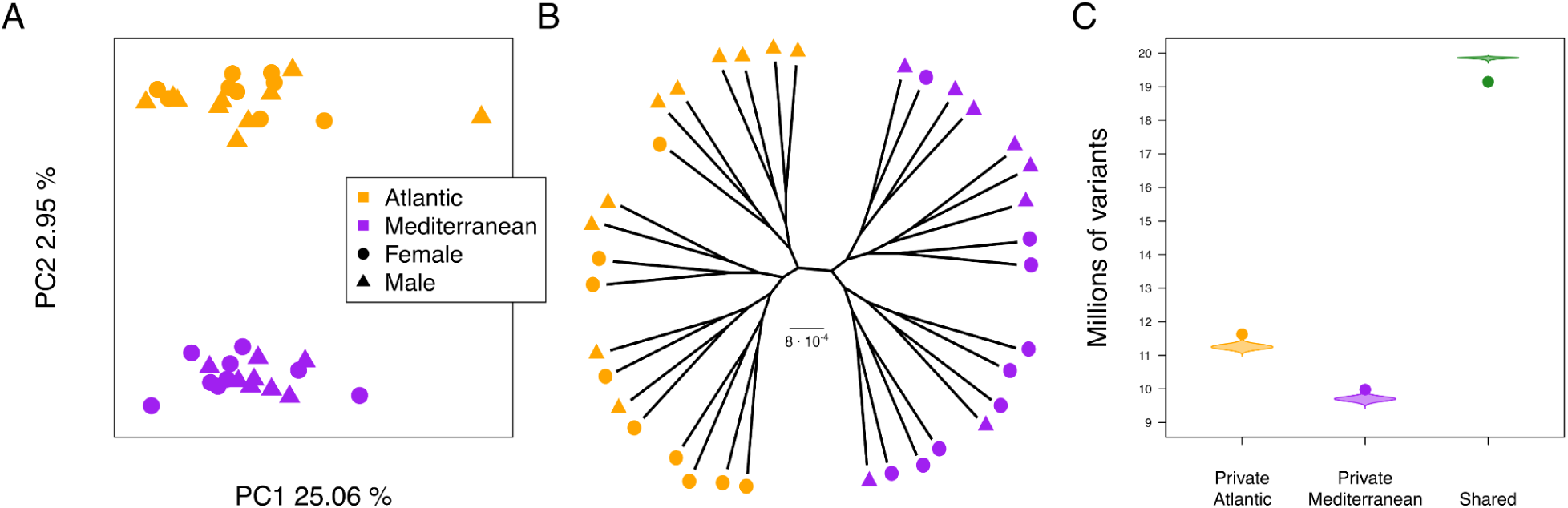
*B. lanceolatum* population differentiation analysis. A) PCA results showing PC1 and PC2. Samples are colored by population and different point shapes indicate different sexes. B) Sample distance tree built with SNPs. Color and point shape as in A. C) Analysis of the number of variants that are segregating in both populations (shared), and the number of variants that are unique to the Atlantic population (private Atlantic) or unique to the Mediterranean population (private Mediterranean). Dots correspond to the observed values. Violin plots represent the expected values under complete absence of population differentiation, which were obtained after reshuffling the labels 10,000 times.

In order to better understand the divergence between specific samples, we built a distance tree (Figure 2B). The resulting tree effectively divides the two populations, indicating that there remains some population differentiation, consistent with PC2. However, the branch dividing the two populations is short compared to the length of the other branches on the tree. In other words, the sum of branch lengths separating two individuals from the same population is just slightly shorter than that separating two individuals from different populations. This aligns with the previous analysis (F_ST_ and PCA) and it shows that the differences between individuals in the same population are much greater than differences between populations. The distance tree also shows the quality of our sampling, illustrating similar distances between all pairs of individuals.

To further investigate population differentiation we ran ADMIXTURE [48]. Given a set number of ancestral populations (K), ADMIXTURE infers the proportion of each individual’s genome that descends from each of the ancestral populations. We tested whether one, two, three or four values of K better fit our data (i.e., had lower cross-validation error); 10 independent iterations were run per K to assess the certainty of the cross-validation estimates. We filtered biallelic SNPs for linkage redundancy using two parallel approaches: LD pruning and distance-based thinning (see Methods). The optimal K value for both linkage redundancy filtering approaches and all iterations, as determined by the lowest cross-validation error, was 1 (Supplementary figure 6). Moreover, K=2 does not clearly separate the two populations. These results suggest that despite a certain level of population differentiation being captured by PC2, our sampling sites belong, in practice, to a single population (K = 1).

In light of our results indicating minimal population differentiation, we conducted a simple statistical test to determine whether the observed level of differentiation was significantly different from that expected in a fully panmictic population (i.e., one with a complete absence of population differentiation). We analyzed the number of variants segregating in both populations (*shared*), as well as those unique to each population (*private*; Figure 2C and Supplementary table 5). 44.93% of the 42,618,268 variants are shared between populations, while 27.29% only segregate in the Atlantic and are fixed in the Mediterranean (private of the Atlantic) and 23.42% are private of the Mediterranean. The remaining variants are either positions that have different fixed alleles in the two populations (*fixed different*, 0.0002%) or positions that are segregating in both populations with different alleles (*variant different*, 4.35%; Supplementary table 5). To have an expectation of these figures in case of complete absence of population differentiation, we performed 10,000 randomizations of the sample location labels and recomputed these statistics in each of them (see Methods, Figure 2C and Supplementary table 5). Note that different sample sizes between populations (19 vs. 17) imply slightly different expectations in terms of private variants. The results support the presence of some population differentiation, yet with a large majority of shared variants. They demonstrate that there are more private variants and fewer shared variants than expected in the complete absence of population substructure, alongside a dominance of shared variants between Atlantic and Mediterranean populations, and an equal prevalence of private variants between the two populations.

Overall, population structure analyses indicate a continuous, substantial, and bidirectional gene flow (i.e., migration) between the Atlantic and Mediterranean sampling sites. Two water currents exist that flow through the Strait of Gibraltar. The first one is superficial and flows west-east, and the other one is deeper and flows east-west [49]. Our results show no disequilibrium between private variants between the Atlantic and Mediterranean sampling sites, suggesting that both water currents are transporting amphioxus embryos and/or larvae.

### Causes of high genomic diversity

In a random mating (i.e., panmictic) population with stable long-term population size, genomic diversity at the genome-wide level is proportional to both N_e_ and μ (θ = 4N_e_μ) [2,22,23]. The *drift-barrier hypothesis* states that high N_e_ and high μ cannot coexist because N_e_ is positively correlated with selection efficiency, and selection will favor low mutation rates given the heavy burden of deleterious mutations [4,50]. Given this context, *B. lanceolatum*’s high whole-genome diversity could be due to (i) high N_e_, (ii) high μ, or (iii) a violation of the random mating assumption. We here perform a series of additional analyses intended to shed light into the causes of *B. lanceolatum* high genomic diversity and support its closeness to panmixia.

The ecology of amphioxus is compatible with approximations to random mating or, in other words, panmixia. First, amphioxus are marine species that reproduce through external fertilization [17–19]. This type of fertilization results in pairings that are close to random, especially compared to pairings resulting from internal fertilization. In external fertilization, natural selection acts only after gamete release, through sperm competition and gamete-level mate choice [51]. In contrast, internal fertilization involves stronger pre-mating sexual selection, resulting in greater deviation from random mating. Second, while adult amphioxus are benthic and have limited mobility, their pre-metamorphic stages are pelagic [17–19]. Before metamorphosis, amphioxus embryos and larvae are planktonic and move with water currents, potentially resulting in migration between local settlements. Finally, these pelagic pre-metamorphic stages are particularly long, lasting from several weeks to months after fertilization, allowing for long-distance migration [17–19].

Supporting amphioxus ecological traits compatible with panmixia, the population differentiation analysis revealed minimal divergence and abundant gene flow between distant sampling sites (Figure 2). This is incompatible with high genomic diversity being caused by pronounced population substructure with low gene flow between isolated populations. To further investigate potential divergence from panmixia, especially introgression that could be the source of high genomic diversity, we analyzed the shape of the *site frequency spectrum* (SFS) and the distribution of the number of heterozygous sites in 50-bp windows. The SFS is heavily influenced by demographic events like introgression, migration or big population size changes. Deviations of the exponential distribution expected under neutrality, constant population size, and no gene flow could be informative of these altering events. Recent introgression, a potential deviation of random mating, would result in a distortion of the SFS [52]. The SFS in *B. lanceolatum* fits the expected exponential distribution and has no deviations, providing no support for introgression as the cause of high genomic diversity (Supplementary figure 4A). The distribution of the number of heterozygous sites in 50-bp windows follows a geometric distribution in the absence of introgression events, and deviations are indicators of such events [13]. In the case of *B. lanceolatum*, the distribution fits perfectly with the expected geometric distribution, which further argues against introgression as the cause of high genomic diversity (Supplementary figure 7). Together, the ecological traits and the absence of both population substructure and recent introgression events strongly suggest that deviations from a panmictic situation are not responsible for the high genomic diversity of *B. lanceolatum*.

The mutation rate per generation per base pair (μ) has recently been estimated for the first time in a close amphioxus species (*B. floridae*) [53]. It was done using deep whole-genome sequencing data of two parents and their offspring. The results estimate a μ in *B. floridae* of 5.89 · 10^-9^ base pairs per generation. This estimate is within the range of what has been estimated in vertebrates [54]. Mutation rates are highly phylogenetically correlated, and significant differences are usually only found at large phylogenetic scales [8,55]. Since *B. lanceolatum* and *B. floridae* diverged between 20 and 150 million years ago [56,57], they are likely to have a similar mutation rate. Another close mutation rate estimate for a more divergent amphioxus species from a different genus (*Asymmetron lucayanum*) corroborates the small variation in mutation rates among amphioxus species [58]. These estimates suggest that the mutation rate is not extremely high in the amphioxus lineage. Therefore, it is unlikely to be responsible for the high genetic diversity observed in several species within this lineage.

Since direct estimates of the mutation rate of *B. lanceolatum* are unavailable, we performed a phylogenetic analysis to further investigate it, taking advantage of the high-quality genome reference and annotation of *B. lanceolatum* [24]. Under neutrality and assuming infinite sites, the mutation rate is equal to the number of substitutions [2,22,23]. Thus we estimated relative substitution rates in amphioxus and other chordates. A phylogenetic tree was constructed using protein-coding one-to-one orthologous genes from two amphioxus species (*B. lanceolatum* and *B. floridae)*, three vertebrate model species (*Homo sapiens, Mus musculus*, and *Danio rerio*), one tunicate species (*Ciona robusta*), one deuterostome species outside chordates (*Strongylocentrotus purpuratus*), and one outgroup species outside deuterostomes (*Drosophila melanogaster*; see Methods; Figure 3). Branch lengths representing average number of substitutions per site are shorter in the amphioxus branch compared to both vertebrate and tunicate species (Figure 3). In other words, the base pair substitution rate in protein-coding genes has been lower in amphioxus than the one in vertebrates and tunicates since their split. Protein-coding genes are genomic regions typically subjected to selection but contain synonymous sites that are largely neutral [22]. Recognizing that the assumptions of neutrality and infinite sites are not fully applicable, we used the number of synonymous substitutions (dS) to make a broad comparison of the mutation rate across the tree (see Methods; Supplementary table 6). The synonymous substitution rate analysis shows a higher dS between *S. purpuratus* and the vertebrate species than between *S. purpuratus* and the amphioxus species (Supplementary table 6). These results roughly indicate a lower (or at least not higher) mutation rate in the amphioxus branch compared to vertebrates and, thus, the absence of a high mutation rate responsible for the high genomic diversity of the amphioxus species.

**Figure 3.**
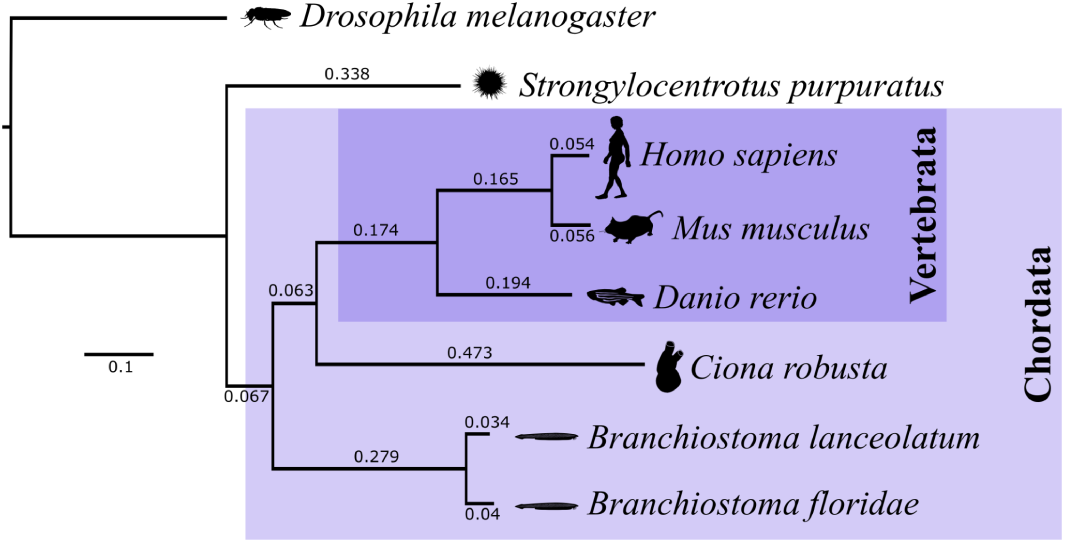
Phylogenetic tree. Phylogenetic tree built with one-to-one protein-coding orthologous genes from *Drosophila melanogaster, Strongylocentrotus purpuratus, Homo sapiens, Mus musculus, Danio rerio*, *Ciona robusta, B. floridae* and *B. lanceolatum*. Branch lengths shown at each branch indicate the average number of substitutions per site. Images from phylopic.org with either Public Domain Mark 1.0 license or CC0 1.0 Universal license.

Finally, we used population genetics simulations to disentangle the effect of high N_e_ from the effect of high μ and compare it with what we observe in the *B. lanceolatum* sequencing data (see Methods). We performed a set of simulations with different combinations of N_e_ and μ values, all resulting in a theoretical genomic diversity (θ = 4N_e_μ)[2] at equilibrium similar to the one estimated in *B. lanceolatum* (Supplementary table 7). In order to compare our data with the simulated genome size (1 kbp), we selected a set of 950 randomly chosen 1-kbp regions of the genome of *B. lanceolatum* (50 per each one of the 19 chromosomes). We tested several statistics and analyses in search of those that would discriminate best between simulations with high N_e_ & low μ or with low N_e_ & high μ. We found that the standard deviation of the heterozygosity and the proportion of singletons and doubletons from all variant sites were the best discriminating statistics. They show that the genomic data of *B. lanceolatum* is closest to the results of the high N_e_ simulations (Figure 4A and 4B). Another discriminative analysis was PCA on the SFS (see Methods; Figure 4C). PC1 (explaining 86.86% of the variance) groups the huge majority of the *B. lanceolatum* genomic windows with the high N_e_ simulations, while high μ simulations show a much more dispersed pattern (Figure 4C). Overall, the comparison of *B. lanceolatum* genomic data with the simulations showed a clear tendency of the genomic data to approach the expectations of high N_e_, further supporting our conclusions.

**Figure 4.**
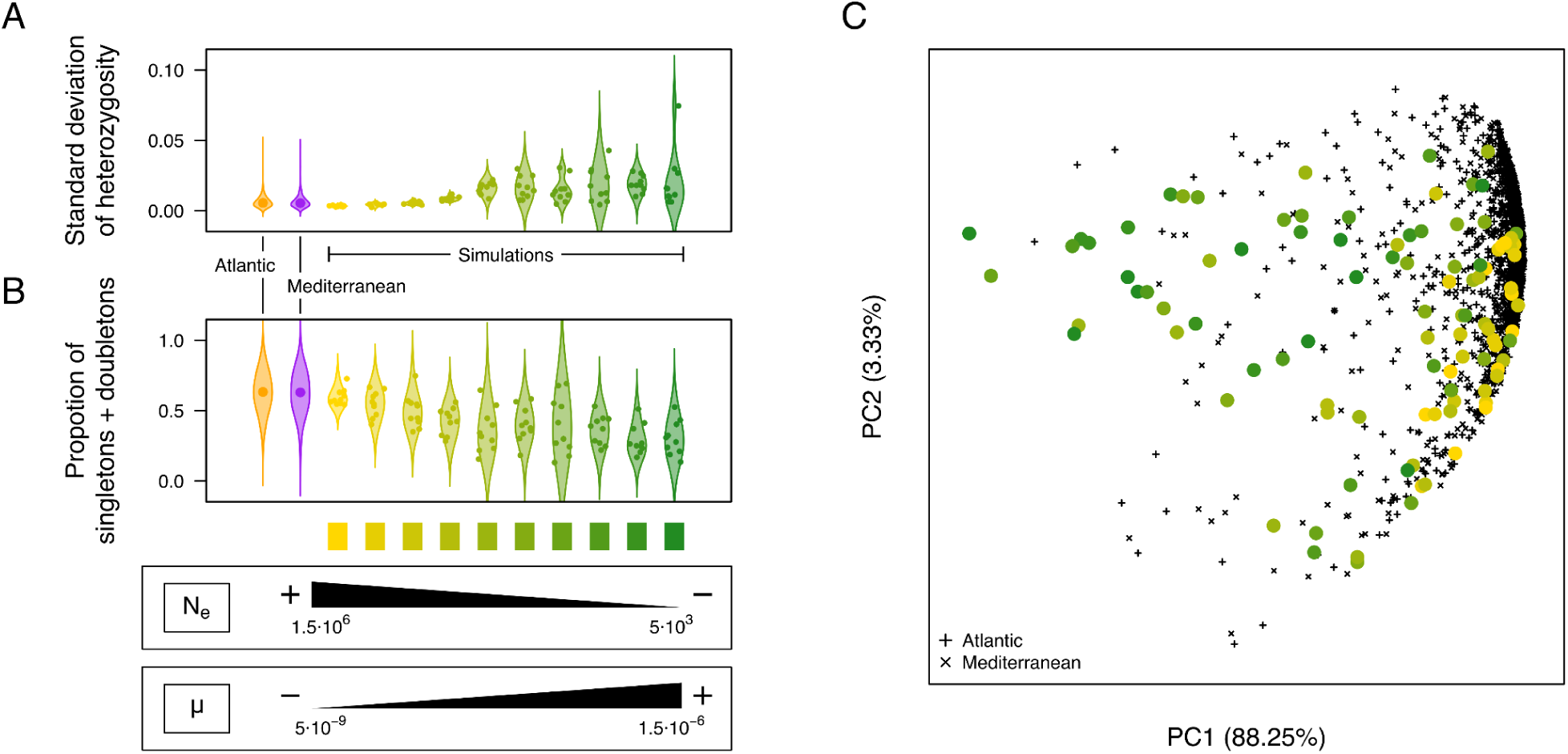
Comparison of *B. lanceolatum* whole-genome data with simulations of molecular evolution. Shades from yellow to dark green depict simulations performed with 10 different combinations of N_e_ and μ that represent a theoretical genetic diversity at equilibrium (θ = 4N_e_μ) [2] similar to the one estimated in *B. lanceolatum*; from yellow depicting high N_e_ (1.5·10^6^ individuals) and low μ (5·10^-9^ base pairs per generation) to dark green depicting low N_e_ (5·10^3^ individuals) and high μ (1.5·10^-6^ base pairs per generation) (see Methods; Supplementary table 7). A) Comparison of the standard deviation of the heterozygosity. *B. lanceolatum* whole-genome sequencing data is represented separately for the Atlantic (orange) and the Mediterranean (purple) populations. For the *B. lanceolatum* sequencing data (orange and purple), violin plots depict the distribution of standard deviation of the heterozygosity in 950 randomly selected 1-kbp genomic regions (50 per each one of the 19 chromosomes), and the point corresponds to its median. For simulated data (yellow to dark green), violin plots depict the distribution of the standard deviation of different simulation runs and points represent the individual simulation run values. B) Comparison of the proportion of singletons and doubletons to the total number of variant sites. Violin plots and points as in A. C) PC1 and PC2 of a PCA performed with the site frequency spectrum of both random 1-kbp windows of the genome of *B. lanceolatum* (vertical crosses for the Atlantic samples and diagonal crosses for the Mediterranean samples) and simulated chromosomes (dots in yellow [high N_e_] to dark green [high μ]).

Using our estimate of per-site nucleotide diversity (π) as θ, and the μ measured in the sister species, *B. floridae* [53], we can estimate the population size of *B. lanceolatum* through the classical relationship θ = 4N_e_μ [2]. This results in a tentative estimate of N_e_ for *B. lanceolatum* of approximately 1.25 million individuals (0.02971 / 4 / 5.89 · 10^-9^ = 1,261,035). Given the similarity of the π estimates for the Atlantic and Mediterranean populations, which suggests similar population sizes, we can take it a step further and estimate the migration rate (m) using an island model with equal subpopulation sizes [60]. Under this model and assumption, m = (1 - F_ST_) / (4N_e_F_ST_) = (1 - 0.00429) / (4 · 1,261,035 · 0.00429) = 4.6 · 10^-5^. The number of migrants per generation is then mN_e_ = 4.6 · 10^-5^ · 1,261,035 = 58 individuals per generation. Please note that these N_e_ and m estimates are highly approximate. They are based on a μ value from a sister species and on an infinite-sites assumption, which is unlikely to be accurate given the species’ genomic diversity (see Discussion).

## Discussion

Amphioxus has been used as a model for evolutionary developmental biology for decades [59]. Indeed, they are the perfect outgroup species to study the evolution of embryonic development of vertebrates [60]. This is because amphioxus are benthic as adults so they are easy to collect, they have a wide worldwide distribution so they can be found in many coastal sampling sites, and, more importantly, they are phenotypically simple [61]. This phenotypic simplicity includes both the fact that their morphology and anatomy are simple and the fact that they are phenotypically very similar between species [19,62]. In this context, before the first amphioxus genome assembly was published, many expected to find amphioxus had a “simple” genome [12]. It turned out not to be the case: amphioxus have a medium size genome (∼500 Mbp), share gene innovation features with vertebrates despite lacking their two rounds of whole genome duplication [24], and have high genomic diversity. The high genomic diversity in amphioxus species has been previously reported in *B. floridae* and *B. belcheri* [12–14]. Here, we thoroughly explore the high genomic diversity of a third species of amphioxus: the European amphioxus (*Branchiostoma lanceolatum*). We provide an unprecedentedly reliable estimate of genomic diversity in an amphioxus species and investigate its causes.

Estimates of genomic diversity are highly dependent on the type of data used and on the methodology employed, and not just on the underlying differences in genomic diversity. The availability and quality of genomic data has increased dramatically during the last two decades. Old diversity estimates performed with few loci or only coding regions can barely be compared to the whole-genome estimates that are now possible. Existing genomic diversity compilations across phyla include estimates obtained using a wide range of data types and methodologies, making comparisons between species quite difficult, especially for highly diverse species [5,7,8,21,35]. In particular, comparing estimates made with data from very few loci, from synonymous sites in protein coding genes, from transcriptomic data, or from only one individual or one population, complicates reliability and interpretation. First, estimates from very few loci can be poorly representative of the whole-genome, especially if they are a specific type of loci (e.g., genic loci, microsatellites) [63]. Second, synonymous sites offer a good neutral comparison against non-synonymous sites, but are by definition within coding sequences in genes (i.e., very often in strong linkage disequilibrium with sites under selection) which have, in general, lower diversity than intergenic regions [22]. Third, since the transcriptome is a functional part of the genome, its diversity is expected to be influenced by selection. Fourth, extrapolating the genomic diversity of a single individual to the whole population depends heavily on the sampled individual’s particular ancestry and can be very biased [45]. Finally, estimates coming from only one population or sampling site in the absence of random mating would be biased by the specific demographic history of that population [6].

The presence of high genomic diversity makes estimation more challenging. In these cases, the sequencing reads differ more from the genome reference, making them more difficult to map accurately. This challenge can be partially overcome by increasing sequencing depth and restricting analysis to more easily mappable parts of the genome. These precautions allow for higher-quality heterozygote and homozygote calls, compensate for random sequencing errors, and reduce incorrectly mapped reads [64]. Furthermore, in highly diverse cases, having a high-quality reference genome is crucial, though it is more challenging to create. The presence of high genomic diversity complicates the distinction between recent paralogs and alternative haplotypes, leading to assembly errors. Incorrectly resolved paralogs and haplotypes in a low-quality reference genome can mislead read mapping and result in biased estimates [65]. Finally, to estimate heterozygosity, one must first determine the extent of the callable length of a genome, given the data and genome reference. Only then can each site be called as either homozygous or heterozygous, and can heterozygosity be calculated. Determining the length of the callable genome in highly diverse cases is especially challenging due to the difficulties on read mapping, distinguishing paralogs, and haplotype resolution.

In the present study, we presented the most reliable estimate of the genomic diversity in an amphioxus species up to date. Previous estimates of genomic diversity in amphioxus were performed with either a single individual or with a single sampling site using a scaffold-level reference genome [12–14]. To obtain a better estimate we took advantage of five factors: whole-genome sequencing data with good coverage; multiple individuals; two distant sampling locations; a high-quality genome reference at the chromosome level [24]; and precise mapping and calling methods. The reference genome of *B. lanceolatum* was built with Hi-C data to properly distinguish paralogy from alternative haplotypes [24]. Our methods were specially designed to accurately identify variant and non-variant sites. To prevent ambiguous variant calling, we used strict quality filters and restricted our analysis to the reliable part of the genome. This included only unique regions with proper sequencing depth coverage for all individuals (i.e., the *callable* genome).

We estimated that approximately 3% of the base pairs in the genome of each *B. lanceolatum* individual are heterozygous. This is a lower-bound estimate, as our analysis was limited to the callable part of the genome where heterozygosity could be accurately determined. The non-callable part of the genome contains a high proportion of repeats and duplications, and it may harbor even more heterozygosity. Nevertheless, 3% of heterozygosity is already a very high proportion of heterozygous sites, and has consequences both at the biological level and when applying common population genomics methods and assumptions to this species. First, extreme levels of genomic diversity, such as those seen in hybrids of diverged species or populations, can result in sterility due to impossibility to resolve recombination events during meiosis [66]. This is clearly not the case for *B. lanceolatum*, as we have shown that its high genomic diversity is due to its large population size and proximity to panmixia. Still, many questions remain unsolved. What are the molecular and cellular consequences, if any, of having such a high genomic diversity due to high population size? Is meiotic recombination or genome 3D structure affected? Are gene expression or its variation impacted? Second, one should take extreme caution when drawing conclusions about species with high genomic diversity using common population genomics methods, as these methods often carry assumptions that may not be applicable in such cases. The main limitation is the frequency of multiallelic variants (i.e., variants having more than two alleles) which occur when a mutation happens in an already variant site. Their existence challenges the common infinite sites assumption, which implies that there are no recurrent mutations. In species with low genomic diversity, there are few multiallelic sites, and they are frequently filtered out or assumed to be so rare that the infinite sites assumption remains valid. Having high genomic diversity increases the probability of multiallelic variants. In the case of *B. lanceolatum*, we observed a non-negligible 11.99% of multiallelic SNPs and 58.03% of multiallelic INDELs, calling into question this assumption. Conversely, another common assumption in population genomics is panmixia. While most studied species are far from panmixia, we demonstrated that *B. lanceolatum* is very close to it. Thus, what is easily assumed in other species (i.e., infinite sites) is difficult to assume in *B. lanceolatum*, while what is difficult to assume in other species (i.e., panmixia) fits *B. lanceolatum* quite well.

Overall, our results confirm that high genomic diversity is very likely to be a common feature of the amphioxus group. In turn, this likely implies that such high genomic diversity has the same underlying cause in all diverse amphioxus species: a high effective population size. Despite needing to assume an underlying infinite sites model to obtain an estimate of effective population size, our results imply extensive gene flow between distant adult local settlements, in this case between the Atlantic and the Mediterranean, that brings *B. lanceolatum* close to panmixia. We estimate that only a small number of migrants per generation between the two subpopulations are needed to maintain this nearly-panmictic context. Since adult amphioxus have low mobility, we deduce that such moderate yet pervasive migration must be achieved through the bidirectional water currents in the Strait of Gibraltar during the embryonic and larval stages, when amphioxus is pelagic [67,68]. Panmixia in *B. lanceolatum*’s wide ecological range results in its large effective population size and its high genomic diversity.

## Methods

### Sampling and sequencing

36 adult *Branchiostoma lanceolatum* individuals were collected from the sandy bottom of the sea at Argelès-sur-Mer, close to the marine station of Banyuls-sur-Mer (European Mediterranean coast), and from the coast of Roscoff (European Atlantic coast). The sampling includes 9 females and 10 males collected in Roscoff, and 9 females and 8 males collected from Banyuls-sur-mer (Figure 1A). The adult European amphioxus individuals were maintained in aquaria during two weeks, dissected in small pieces and then flash frozen in liquid nitrogen. DNA was extracted using the DNeasy blood and tissue kit from QIAGEN, following the manufacturer’s instructions. The samples were sent for paired-end whole-genome sequencing at the Genomic Technologies Facility of the University of Lausanne, where they were sequenced in two different batches. The first batch was performed in December 2020 with an Illumina HiSeq4000 sequencer. The second batch was performed in order to increase sequencing coverage in April 2023 with an Illumina NovaSeq6000 sequencer (Supplementary table 1). Each sample had three different fastq file pairs coming from three different lanes, one from the first batch and the other two from the second.

### Variant calling

We used *BWA MEM2* (version 0.7.17) [25] to map each pair of fastq files (per sample and lane) to the high quality reference genome of *B. lanceolatum*, BraLan3 [24]. The high quality of this reference genome is essential for our analysis, although it restricts our analysis to the nuclear chromosomes since the mitochondrial chromosome is missing from the reference. We then followed the best practices recommendations of GATK (version 4.2.4.0) [26] to call small variants (SNPs and INDELs) on non-model organisms (Supplementary figure 1) [26]. We did it by chromosome for computational optimization purposes. This procedure included two rounds of base recalibration that were applied as described below. After sorting each sample-lane sam file and converting it to a bam file with *samtools* (version 1.17) [69] we split it by chromosome resulting in sample-lane-chromosome bam files. We then merged the three bam files corresponding to different lanes for each sample and chromosome obtaining a single bam file per sample and chromosome. We did so while marking duplicated reads with *samtools* for *MarkDuplicatesSpark* (GATK4 version 4.2.4.0). Then, we did a first variant calling per sample and chromosome using *HaplotypeCaller*. We merged all samples’ calls (VCF files) for each chromosome, obtaining one single VCF file per chromosome (*GenomicsDBImport* and *GenotypeGVCFs*). We subjected each chromosome’s calls to a hard filtering following GATK best practices recommendations. We split by variant type (SNPs and INDELs) using *SelectVariants*, applied the recommended hard filters for SNPs (QD < 2.0; QUAL < 30.0; SOR > 3.0; FS > 60.0; MQ < 40.0; MQRankSum < -12.5; and ReadPosRankSum < -8.0) and for INDELs (QD < 2.0; QUAL < 30.0; FS > 200; and ReadPosRankSum < -20.0) and merged them back using *MergeVcfs*.

After this first variant calling and hard filtering, we proceeded to the first round of base recalibration using the first per chromosome hard filtered calls as the set of known sites. We performed base recalibration on each sample-lane-chromosome bam file (*BaseRecalibrator*) and applied it to obtain the new base-recalibrated sample-lane-chromosome bam (*ApplyBQSR*). We then repeated the procedure described above in the same exact way; merged all lanes for each sample and chromosome, performed a variant calling per chromosome and sample, merged all sample calls for each chromosome and hard-filtered them. With this we obtained a second set of hard-filtered calls per chromosome that we used as known sites for the second round of base recalibration conducted in the exact same way as the first. After the second base recalibration, we repeated the merging, variant calling and hard filtering per chromosome to obtain a third set of hard-filtered calls that we then subjected to several additional filters described in the following section.

### Additional variant filters

After following the GATK best practices recommendations for non-model organisms to obtain a set of small variants (SNPs and INDELs), we applied several additional filters (Supplementary figure 1; Supplementary table 2). First, we applied a mappability masking only considering variants within *SNPable regions* c=3 following the procedure in http://lh3lh3.users.sourceforge.net/snpable.shtml (source code downloaded the 11th of April 2023). Briefly, this method splits the genome reference in 35 bp segments that afterward maps back to the genome and calibrates the quality of the resulting mapping in each genomic position. High level SNPable regions (c=3) include only positions for which most of the overlapping 35-mers are mapped uniquely and without mismatches. This mappability masking was used to ensure we are only considering variants in unique regions (non-repetitive, non-duplicated) with ensured high quality mapping. Second, we only considered variants in sites with a minimum depth of coverage among all samples of >= 5 and < 20. In this second filter, we ensure all samples have enough coverage to distinguish a homozygous site from a heterozygous site in all considered sites. This filter excludes duplicated regions that avoided the mappability filter due to their absence in the reference (i.e., copy-number variants). Third, we discarded very isolated sites that passed the two first filters by only considering variants with >= 20% of the sites passing the two first filters in a sliding window approach with window size of 100 kbp and a step of 10 kbp. These first three filters restrict our analysis to the part of the genome that is unique and for which we can reliably call both homozygous and heterozygous variants for all individuals, the *callable genome*. Fourth, we considered only variants present in our sample by excluding variants private to the reference (AF == 1.00 in our sample with *VariantFiltration*). Fifth, we filtered out the variants with missing genotype in any of the samples (*vcftools --recode --max-missing 1*; version 0.1.16). The resulting list of variants, both SNPs and INDELs, was used for the downstream analysis.

### Genome mapping and variant calling method comparison

We compared two genome mapping tools (BWA MEM2, which was used in the main analysis, and Minimap2 [27]) and two variant callers (GATK, which was used in the main analysis, and FreeBayes [28]). In total, four different mapper-variant caller combinations of methods were compared (Supplementary figure 3). This comparison was run on the four smallest chromosomes of BraLan3 (chromosomes 16, 17, 18 and 19). Minimap2 (version 2.30) was run with the following arguments: *minimap2 -ax sr -B4 -O5,24 -E2,1*. FreeBayes (version 1.3.10) was run with the following arguments: *freebayes --haplotype-length 0 --use-best-n-alleles 10 –theta 0.02 --min-alternate-fraction 0.05 --min-alternate-count 2 --min-base-quality 10 --min-mapping-quality 20 --genotype-qualities*. Additional variant filters were applied after each mapper-variant caller method as in the main analysis.

### Genomic diversity analysis

We used *VariantsToTable* to extract genotypes from the VCF file. Then we used in-house scripts (see Data and code availability) to calculate heterozygosity per sample, nucleotide diversity (π), and the number of alleles per variant. We calculated heterozygosity per sample by counting the number of heterozygous sites in a given region of the genome and dividing it by the length of that region. Total heterozygosity for a given sample, for example, is equal to the total number of heterozygous sites of that sample in the callable genome divided by the length of the callable genome. Nucleotide diversity, or the average pairwise differences, was calculated for a group of samples (either all Atlantic, all Mediterranean, or all samples) and normalized by sequence length. We used the following formula:

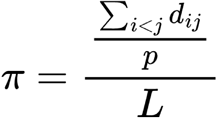

*L* corresponds to the length of the region in question, *p* corresponds to the number of pairwise comparisons and *d_ij_* corresponds to the number of nucleotide differences between sequences *i* and *j* in the region in question [29]. Using the gene annotation from BraLan3, we divided the genome into exons, introns, promoters (1 kbp upstream of all genes), and intergenic regions [24]. We then evaluated heterozygosity and nucleotide diversity in the total length of the callable genome, as well as in each of these groups of regions.

### Population structure

We calculated the population F_ST_ per site with *vcftools* directly from the VCF file with the option *--weir-fst-pop* per population and averaged all variant values to get the total population F_ST_.

To include multiallelic variants in the Principal Component Analysis (PCA), we built a matrix in which columns were samples and rows were alleles (instead of variants). Each cell contained 0, 1 or 2 depending on the number of copies of the corresponding allele carried by the corresponding sample. With this matrix, we used the R package ade4 (version 1.7.22) to perform PCA.

To build the sample distance tree, we built a consensus genomic sequence for each sample. In order to avoid misalignments, we only considered SNPs (not INDELs) to build the consensus sequences (*vcftools –recode --remove-indels* and *bcftools view -e ‘ALT=“*”’*). We used *bcftools consensus* (version 1.17) with the *-I* option (IUPAC codes) to have the diploid consensus sequence in a haploid-like format. We then joined all the sample consensus genomic sequences and used *iqtree* (version 2.3.0) to build the distance tree and *FigTree* (version 1.4.4) to visualize it.

To get an expected distribution of values for the number of variants shared between populations, private to each population, fixed different, and variant different, we shuffled the population affiliation labels 10,000 times and recalculated each value every time. With this, we got a distribution of expected values for each one of these types of variants in the absence of population substructure. To test how extreme the observed values are from the distribution of values expected under no substructure, we performed an empirical permutation one-sided test.

We ran ADMIXTURE (version 1.3.0) [48] on biallelic SNPs to determine which of the K values (1, 2, 3 or 4) was optimal for our data. We selected and filtered biallelic SNPs having a minimum threshold of depth (DP) of 5, genotype quality (GQ) of 20, site quality (QUAL) of 30 and allowing a maximum of 20% of missing genotypes across samples (F_MISSING). We did so using *bcftools* (version 1.21; *bcftools filter -e ‘FORMAT/DP < 5 M FORMAT/GQ < 20’ --set-GTs . ${VCF} | bcftools view -e ‘F_MISSING > 0.8’ --types snps -O z -o ${filteredSNPsVCF}*) and *vcftools* (*vcftools --gzvcf ${filteredSNPsVCF} --remove-filtered-all --minQ 30 --min-alleles 2 --max-alleles 2 --recode --recode-INFO-all --out ${filteredbialelicSNPsVCF}*). We kept 29,758,921 biallelic SNPs across all chromosomes. We then performed two parallel SNP redundancy filtering strategies. First, we performed an LD pruning using *plink2* (version 2.00a4.3) [70] with a window size of 50 kbp, a step size of 10 and a r^2^ threshold of 0.3 (*plink2 --vcf ${filteredbialelicSNPsVCF} --bad-ld --double-id --allow-extra-chr --set-missing-var-ids @:# --indep-pairwise 50 10 0.3 --out ${toprune}; plink2 --vcf ${filteredbialelicSNPsVCF} --double-id --allow-extra-chr --set-missing-var-ids @:# --extract ${toprune}.prune.in --recode vcf --out ${pruned}*). After *plink2* LD pruning, 10,764,841 SNPs remained out of a possible 29,758,921 biallelic SNPs. Given that LD pruning is not optimal in a sample size of less than 50 individuals, we performed an alternative approach accounting only for physical linkage. This alternative approach was thinning using *vcftools* that removes redundant SNPs by distance (*vcftools --gzvcf ${filteredbialelicSNPsVCF} --thin 500 --recode --recode-INFO-all --out ${thinned}*). This second method retained 557,118 SNPs out of a possible 29,758,921 biallelic SNPs. For each SNP redundancy filtering approach, we then merged all chromosome VCF files. We converted the concatenated VCF files to plink format files using vcftools and plink2. We ran 10 independent iterations of ADMIXTURE for each K value, after each of the two SNP redundancy filtering approaches. In total, this equated to 80 runs of ADMIXTURE. We used a different seed for each run to ensure independence between runs. Individual genomic proportions between populations and cross-validation errors per K were consistent across iterations and between the two linkage-filtering approaches. K=1 presents the lowest cross-validation error in all 10 iterations, consistently for both SNP redundancy filtering approaches.

### Phylogenetic analysis

We downloaded protein sequences from Ensembl (release 111) for *Homo sapiens* (GRCh38), *Mus musculus* (GRCm39), *Danio rerio* (GRCz11), *Drosophila melanogaster* (BDGP6.46) and *Ciona robusta* (KH), from Ensembl Metazoa (release 58) for *Strongylocentrotus purpuratus* (Spur_5.0) and from NCBI genome database for *Branchiostoma floridae* (Bfl_VNyyK). For *B. lanceolatum* we used the gene annotation and protein sequences of BraLan3 [24]. We selected the longest protein sequence per gene when more than one was available in each case. With all the protein sequences of all species we build orthogroups with *OrthoLoger* (version 3.0.5 using *brhclus* 5.1.7, *blast* 2.14.1, *swipe* 2.1.1, *diamond* 2.1.8, *mmseqs2* 15.6f452, *cdhit* 4.6 and *expect* 5.45.4) [71]. Only the one-to-one orthogroups that were shared by all species were used in the following analysis.

To build the phylogenetic tree, we build a multiple sequence alignment (MSA) of each orthogroup protein sequence using *MAFFT L-INS-i* (version 7.526) [72]. We trimmed each alignment with *ClipKIT* (version 2.3.0) [73]. We then concatenated the protein MSAs of all orthogroups and used the resulting concatenated MSA to build the phylogenetic tree with *IQ-TREE* (version 2.3.0) using the GTR20 model and 1,000 bootstrap replicates [74]. To estimate the rate of synonymous substitutions along the phylogenetic tree, we back-translated each orthogroup protein MSA to the corresponding transcript sequences obtaining one transcript MSA per orthogroup using *TreeBeST backtrans* (version 1.9.2_ep78) . We trimmed the resulting transcript MSAs using *ClipKIT* and concatenated them. We used the resulting concatenated MSA as well as the previously built phylogenetic tree as inputs for *CODEML* (*PAML* version 4.10.7), which we ran with seqtype = 1, CodonFreq = 2, clock = 0, aaDist = 0, model = 0, NSsites = 0, icode = 0, Mgene = 0, fix_kappa = 0, kappa = 2, fix_omega = 0, omega = .4, fix_alpha = 1, alpha = 0, Malpha = 0, ncatG = 3, fix_rho = 1, rho = 0, RateAncestor = 0, Small_Diff = .5e-6, cleandata = 0, fix_blength = 0, method = 0.

### Simulations

Using SLiM (version 4.1) [75], we performed simulations of a single population’s neutral evolution with 10 different combinations of constant N_e_ and μ, all resulting in a theoretical θ of 0.03, similar to the one measured in *B. lanceolatum*. N_e_ ranged from 5,000 to 1,500,000 individuals, while μ ranged from 1.5·10^-6^ to 5·10^-9^mutations per base and generation (Supplementary table 7). We performed 10 replicas of each N_e_/μ combination that were run with a nucleotide base mode (*initializeSLiMOptions(nucleotideBased=T)*) to allow for multiallelic sites, a chromosomal size of 1,000 nucleotides, neutral mutations with a 0.5 dominance coefficient and a Jukes–Cantor (JC69) nucleotide substitution model (*initializeMutationTypeNuc(“m1”, 0.5, “f”, 0.0), initializeGenomicElementType(“g1”, m1, 1.0, mmJukesCantor(μ/3))*) and a constant recombination rate of 1·10^-8^ (*initializeRecombinationRate(1e-8)*). All the simulations were run for 3,000,000 generations.

## Data and code availability

All the genomic whole-genome sequencing raw data (FASTQ files) is available in the European Nucleotide Archive (ENA) under the project PRJEB115533. The variant calls of our study (VCF files) are available in the European Variation Archive (EVA) under the project PRJEB120745 and the analysis ERZ29712263. All the code from raw data to figures is built in a snakemake pipeline to allow for reproducibility and can be found in https://github.com/marinabraso/AGenDiv, Admixture analysis can be found in https://github.com/Angelica-Pulido/Amphioxus_panmixia. All the code can be found in a frozen version in Zenodo (https://doi.org/10.5281/zenodo.21503009).

## Acknowledgements

We thank Sagane Joye and Giulia Campli for sharing code and expertise to run OrthoLoger. We also thank Jérôme Goudet, Tristan Cumer, and other members of DEE for useful feedback. Research by M.B.-V. and M.R.-R. is funded by the Swiss National Science Foundation grant 207853. This work benefited from access to the Observatoire Océanologique de Banyuls-sur-Mer, an EMBRC-France and EMBRC-ERIC site. The laboratory of H.E. and S.B. was supported by the CNRS, and by the “Agence Nationale de la Recherche” under the grants ANR-21-CE13-0034. S.B. is supported by the Institut Universitaire de France (IUF). The funders had no role in study design, data collection and analysis, decision to publish, or preparation of the manuscript.

## Supplementary figures

**Supplementary figure 1.**
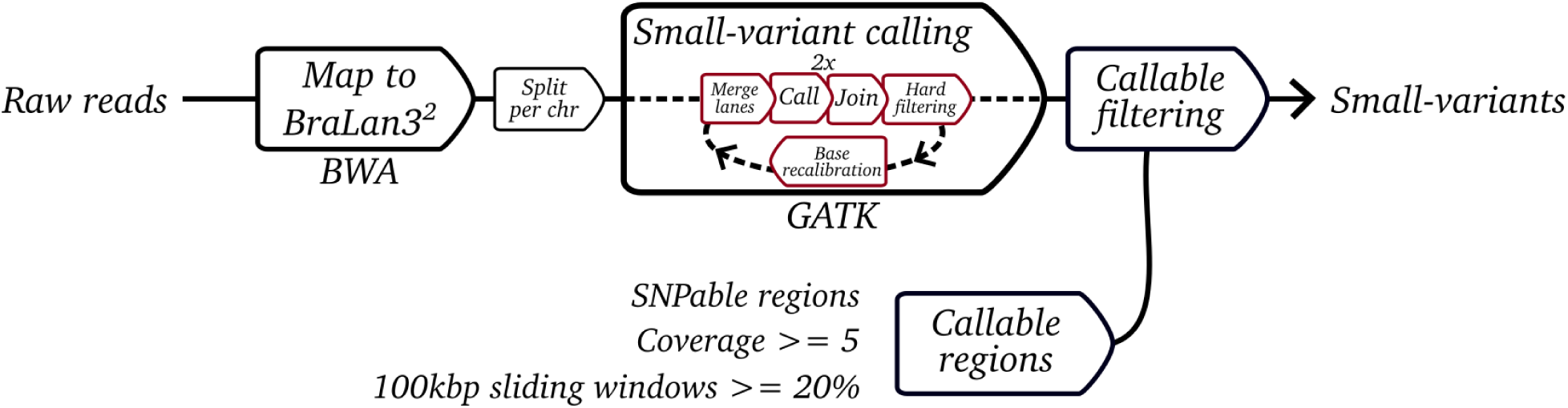
Small variant (SNPs and INDELs) calling methodology.

**Supplementary figure 2.**
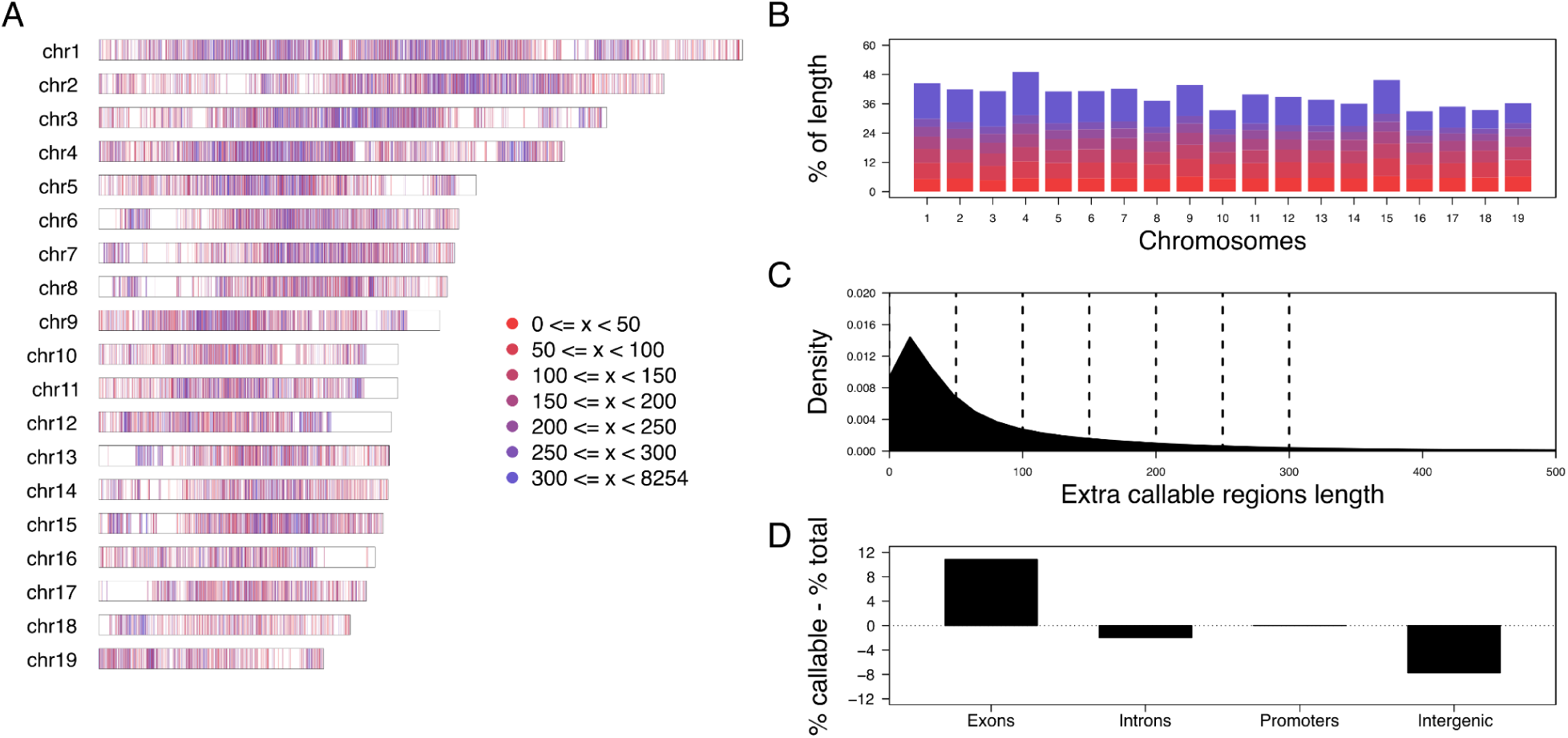
Description of callable regions. A) Distribution of callable regions along each chromosome. Regions are colored according to their length. B) The percentage of chromosome length covered by callable regions of different sizes. Colors as in A. C) Length distribution of callable regions. D) The difference between the percentage of callable length and genome length covered by each feature (Supplementary table 3). All four cases are statistically different from a random distribution of callable regions in a Fisher’s exact test (Supplementary table 3).

**Supplementary figure 3.**
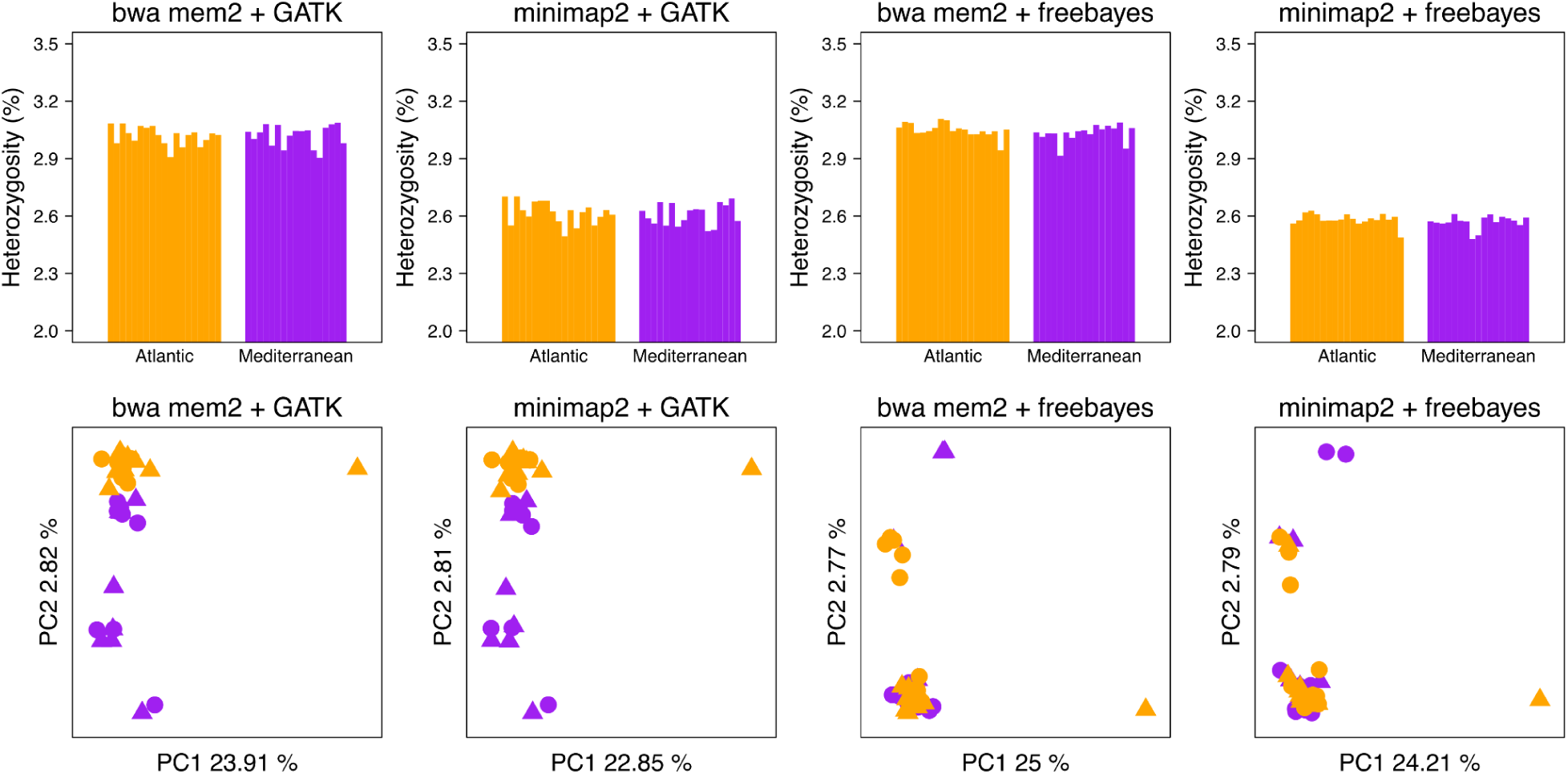
Comparison of genome mapping and variant calling methods. This comparison includes chromosomes 16, 17, 18 and 19. We used two genome mappers (BWA MEM2 and Minimap2) combined with two variant callers (GATK and FreeBayes). In total there are four method comparisons (from left to right; BWA MEM2 + GATK, Minimap2 + GATK, BWA MEM2 + FreeBayes, Minimap2 + FreeBayes). Since they are central to the overall result of this study and its interpretation, we reproduced Figure 1B (total heterozygosity per sample, top) and Figure 2A (PCA, bottom) for all four combinations of methods. Color and point shape as in Figure 2A. Overall heterozygosity is found to be robust to variant calling methods, but slightly affected by mapping methods. Conversely, PCA is robust to different mappers but sensitive to variant calling methods, though this does not affect the interpretation of the overall result. In summary, these results demonstrate the robustness of our methods, showing that slight changes in specific numbers resulting from different methods do not affect the interpretation of the overall result.

**Supplementary figure 4.**
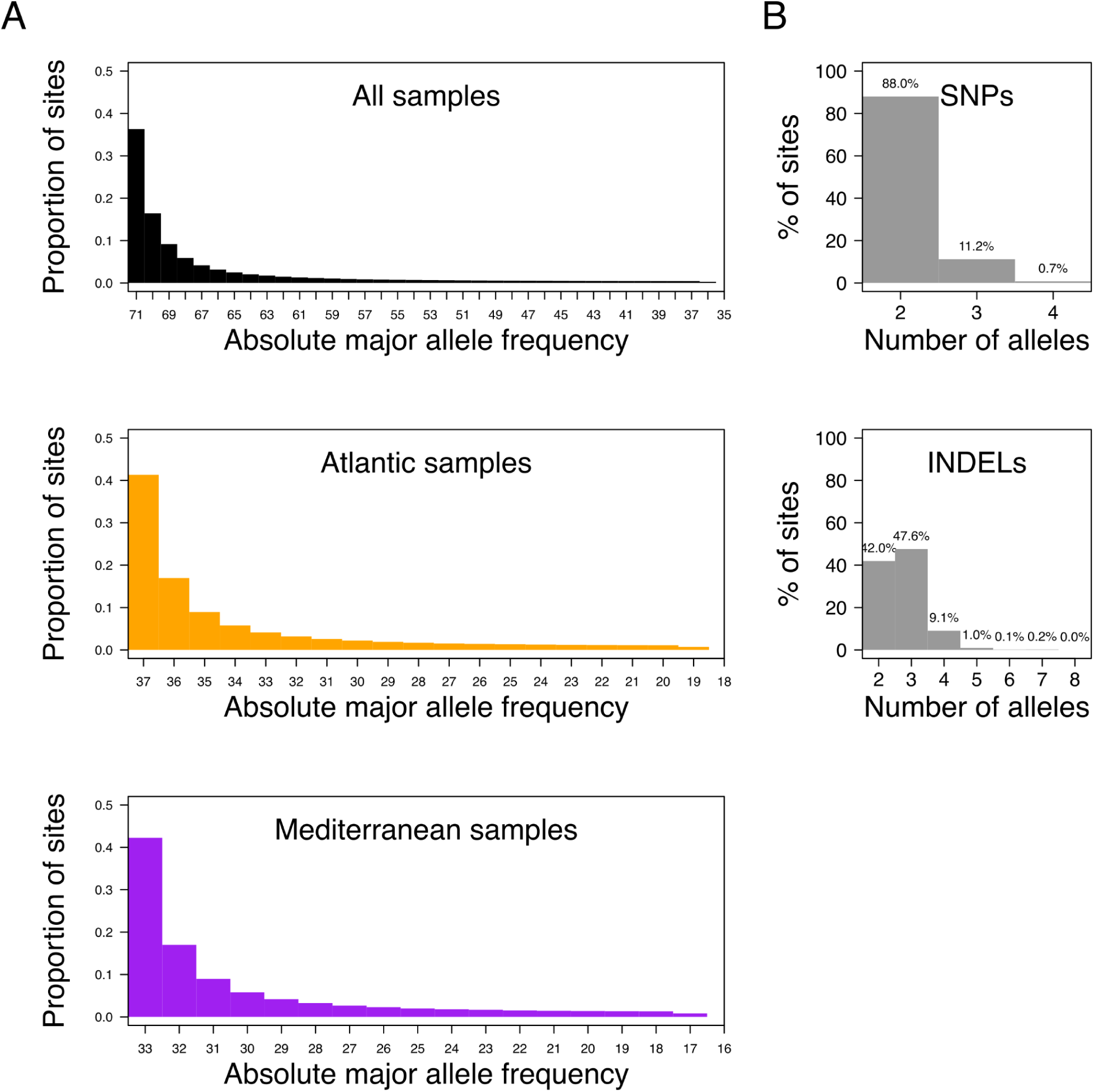
SNP and INDEL site frequency spectrum and distribution of allele number. A) Site frequency spectrum (SFS) of both SNPs and INDELs for all samples (top, black), Atlantic samples (middle, orange) and Mediterranean samples (bottom, purple). B) Distribution of the number of alleles per variant for SNPs (top) and INDELs (bottom).

**Supplementary figure 5.**
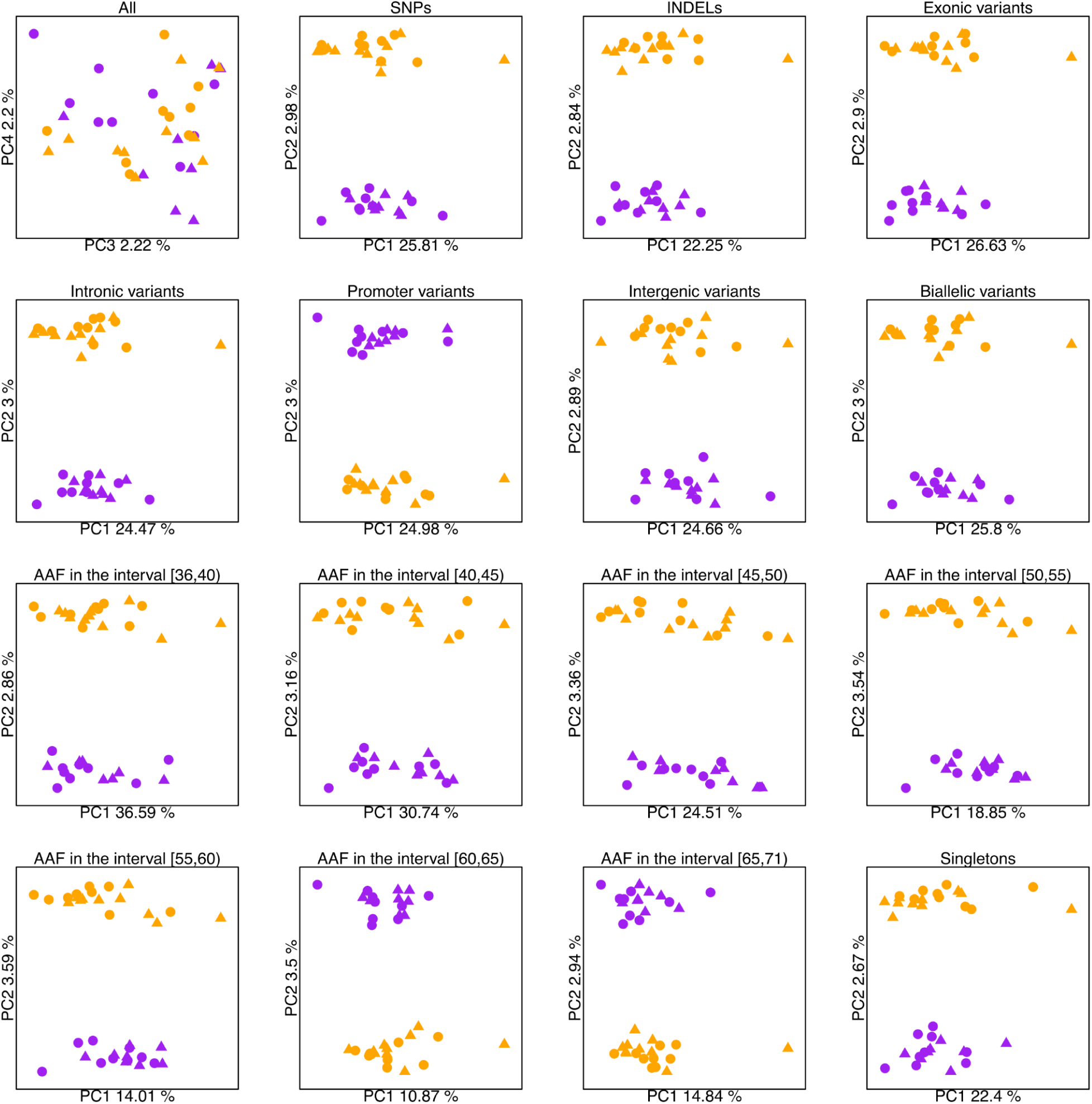
PCA on different subsets of variants. AAF refers to absolute allelic frequency. Color and point shape as in Figure 2A.

**Supplementary figure 6.**
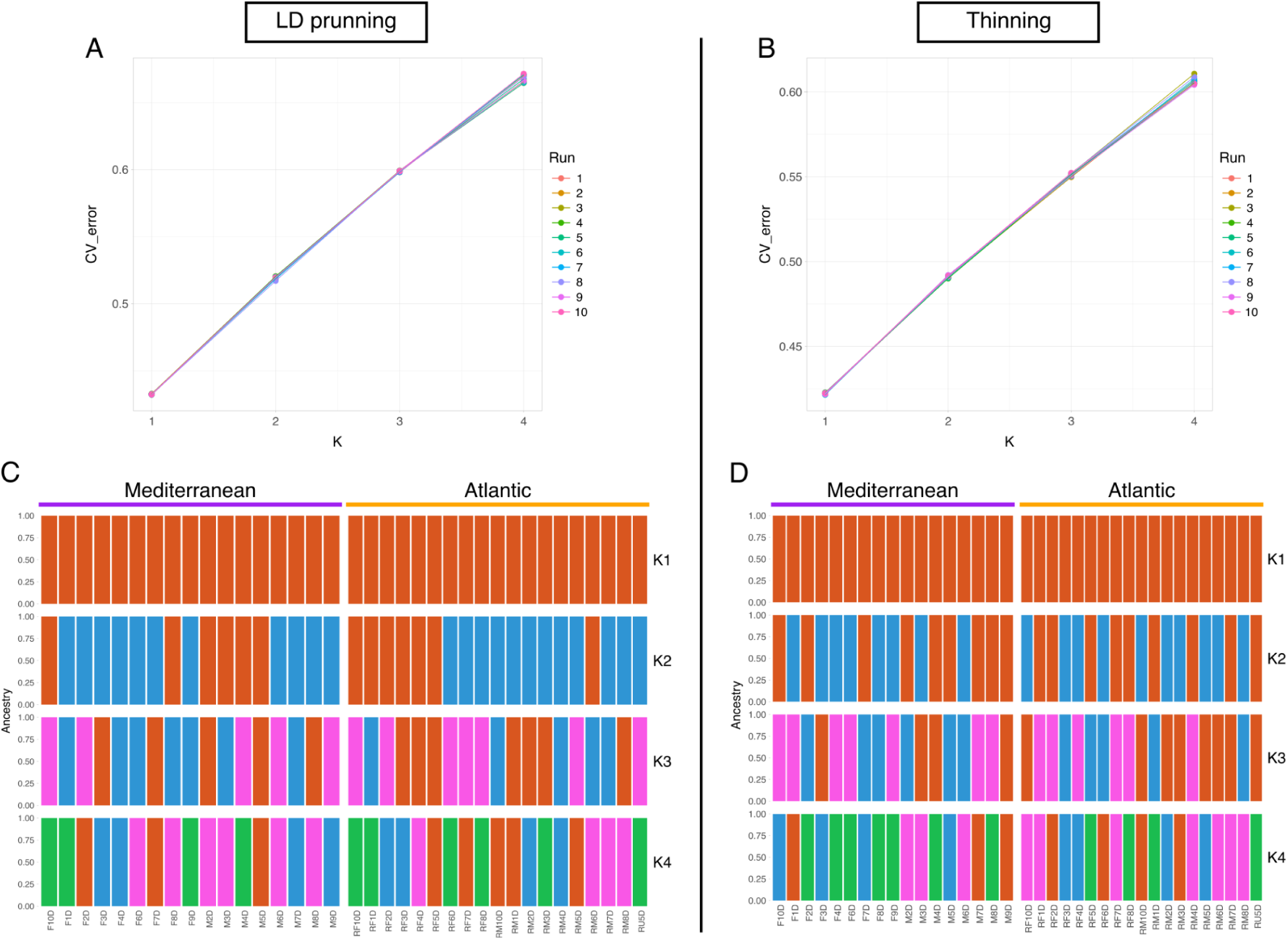
ADMIXTURE analysis results. A and C correspond to the LD pruning approach while B and D correspond to the distance thinning approach. A and B show the cross-validation error for each tested K value, for 10 iterations per K. C and D show the individual genomic proportions that descend from the inferred ancestral population for each test K value at iteration number 2. Note that, in all cases, the proportion of the genome belonging to one of the inferred ancestral populations is sufficiently high for the others to be unappreciable in the figure. Further details on these results can be found in the corresponding GitHub repository (see Methods).

**Supplementary figure 7.**
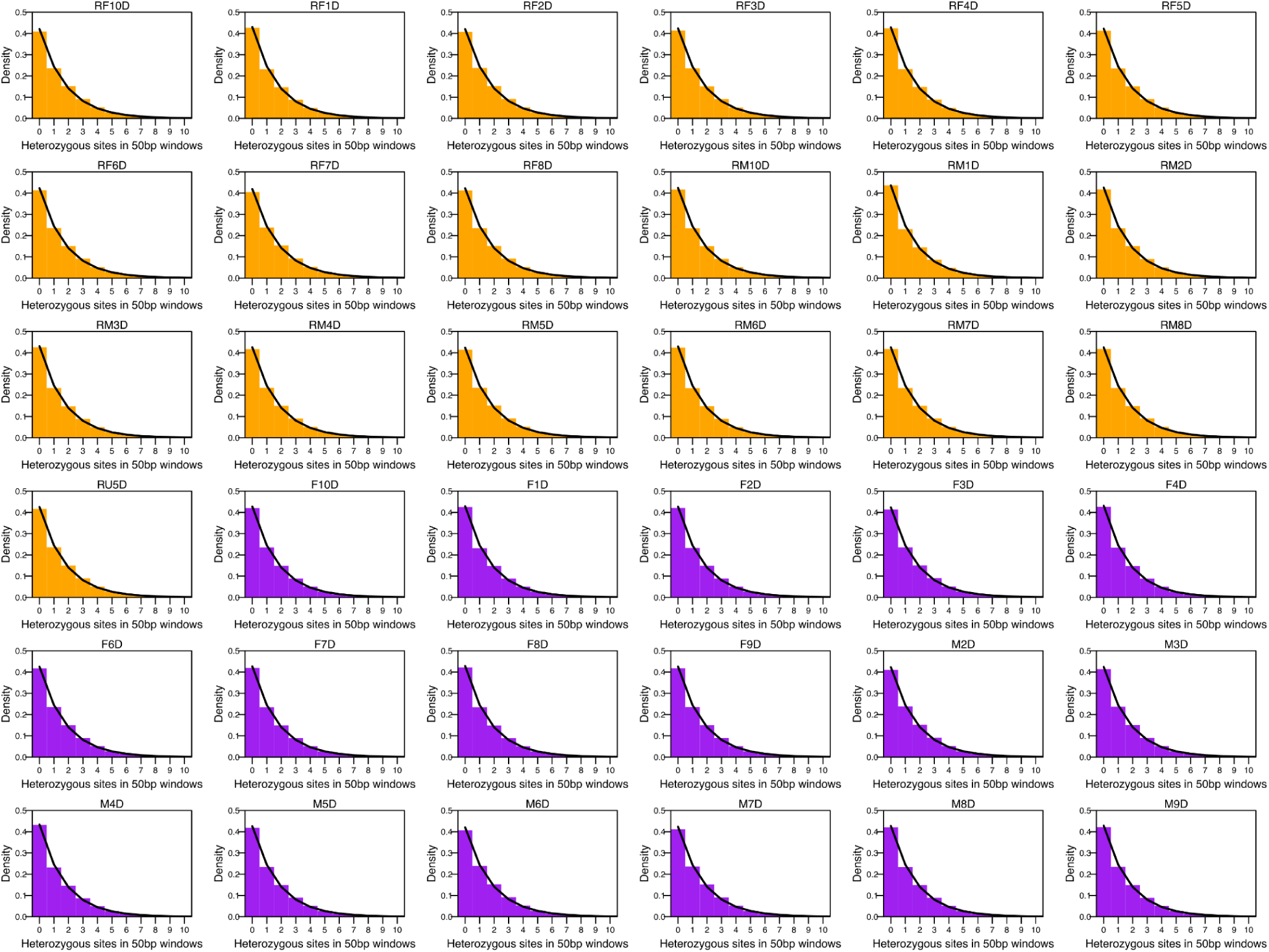
Geometric distribution of heterozygous sites per window. Distribution of the number of heterozygous sites in 50-bp windows fitted to a geometric distribution for Atlantic samples (orange) and Mediterranean samples (purple).

## Notes

https://doi.org/10.5281/zenodo.21503009

### Competing Interest Statement

The authors have declared no competing interest.

### Summary of Updates

We revised the refence list and detected some typesetting errors. We have now corrected them.

